# Psychophysiological Evidence for Fear Extinction Learning via Mental Imagery

**DOI:** 10.1101/2021.06.17.448826

**Authors:** Xinrui Jiang, Steven Greening

## Abstract

Imagery-based extinction procedures have long been used in the treatments of fear-related conditions. The assumption is that imagery can substitute for the perceptual stimuli in the extinction process. Yet, experimental validations of this assumption have been limited in number and some have relied exclusively on measures of autonomic reactivity without consideration of conscious feelings of fear. The current investigation sought to assess whether imagery-based exposure could lead to extinction of conditioned fear to the corresponding perceptual stimulus. Conditioned fear responses were measured by both a physiological (i.e., skin conductance response/SCR) and a subjective (i.e., self-reported fear) measure. Participants (*N* = 56) first underwent perceptual differential fear conditioning, then imagery extinction, then perceptual extinction. SCR evidence was found for successful fear conditioning, generalization of fear from viewing to imagery, and most importantly, the absence of differential fear after imagery extinction upon re-exposure to the conditioned perceptual stimulus. Self-reported fear confirmed the acquisition and generalization of fear and provided evidence of a significant reduction in differential fear conditioning across extinction. Consistent with clinical evidence of the efficacy of imagery extinction and the existing limited experimental literature, the current study offers support for fear extinction to perceptual stimuli via imagery exposure.

## 1. Introduction

From as early as the practices of shamanism some 20,000 years ago (Achterberg, 2002) to today’s imagery-based cognitive behavioral therapy techniques (Holmes, Arntz, & Smucker, 2007), mental imagery has been harnessed for its efficacy as a cognitive process that affects emotion perception and regulation. It is commonly referred to as a perceptual-like experience in the absence of external sensory input (Kosslyn, Thompson, & Ganis, 2006). Imagery is recognized as an important component of various fear-related affective conditions in terms of both their symptoms (e.g., intrusive images in depression, flashbacks in post-traumatic stress disorder or PTSD; Matthews, Collins, Thakkar, & Park, 2014; Weßlau & Steil, 2014), and clinical treatments (e.g., imaginal exposure, Abramowitz, 1996; Hecker, 1990; Powers, Halpern, Ferenschak, Gillihan, & Foa, 2010; Turner, Beidel, & Jacob, 1994; imagery rescripting, Arntz, 2012). The primary explanation for imagery’s impact in these examples is that it can substitute for the actual stimuli in affective processes to regulate fear responses. Yet, only a few experimental examinations have been done to assess this explanation (Mertens, Krypotos, & Engelhard, 2020).

While some have suggested that informational processing attributed to mental imagery is purely propositional, an abundance of behavioral and brain imaging research suggests that mental imagery is at least partially depictive or pictorial (Kosslyn, 1981, 2005; Kosslyn, Ganis, & Thompson, 2001). According the depictive theory, mental imagery generates a neural representation that functions like a weaker version of perception and is associated with the subjective experience of, for example, “seeing with the mind’s eye” (Kosslyn et al., 2001). Thus, imagery of an object can “stand in” for the corresponding external stimulus in various cognitive processes including associative learning and other affective processes (Lewis, O’Reilly, Khuu, & Pearson, 2013; Pearson & Kosslyn, 2015; Pearson, Naselaris, Holmes, & Kosslyn, 2015). On the other hand, the bio-informational theory noted that mental imagery contains not only the perceptual information of the imagined stimulus but also its conceptual information stored as propositions such as “relationships, descriptions, interpretations, labels, and tags” (Lang, 1977, 1979). The bio-informational theory posits that it is through the activation of this propositional network during imagery that emotional modifications occur (Lang, 1977, 1979). Regardless of the precise nature of mental imagery, it is acknowledged by both theoretical perspectives that mental imagery has the capacity to elicit emotional characteristics similar to its perceptual counterparts.

In addition to successful clinical applications and theoretical foundations, neuroimaging research has also provided evidence that mental imagery contributes to the elicitation or modification of emotional reactivity similarly to perception. For example, meta-analytic reviews of the brain regions associated with the emotional processing of perceptual stimuli report activations in regions such as medial prefrontal cortex (mPFC), anterior cingulate cortex (ACC), insula (INS), and amygdala (Kober et al., 2008; Murphy, Nimmo-Smith, & Lawrence, 2003; Phan, Wager, Taylor, & Liberzon, 2002). Importantly, activations of the same regions have been reported during elicitation of emotions via mental imagery, including mPFC (Schaefer et al., 2003), ACC, INS (Phan et al., 2002), and amygdala (Kreiman, Koch, & Fried, 2000). Taken together, the existing research appears to support the intuition that mental imagery of an emotional stimulus can substitute for its external presentation. In contrast to the findings noted above, relatively less experimental research has investigated the emotion regulatory capacity of mental imagery as compared to perception. Such a comparison is a direct way to investigate the assumed correspondence between mental imagery and perception.

Among the existing studies that investigated the connection between mental imagery and emotion, many based their task paradigms on differential fear conditioning (Agren, Björkstrand, & Fredrikson, 2017; Greening et al., 2021; Grégoire & Greening, 2019; Koizumi et al., 2016; Reddan, Wager, & Schiller, 2018). Such paradigms typically involve one initially neutral conditioned stimulus (CS+) which is repeatedly paired with an aversive or threatening unconditioned stimulus (US; e.g., a loud noise or mild electric shock), and a second CS (CS-) which is never accompanied by the US (Carter, Hofstotter, Tsuchiya, & Koch, 2003). Upon such pairing, the CS+ but not the CS- exhibits a conditioned response (CR) that is similar to the type of response elicited by US (e.g., increased self-reported fear; increased skin conductance response, SCR; increased heart rate; or, escape behavior; Carter et al., 2003; Critchley, Mathias, & Dolan, 2002). The acquisition of differential fear conditioning is quantified as the difference in the CR between the CS+ versus the CS-. Greening et al. (2021) observed recently that visual imagery of conditioned stimuli elicited a significant differential fear conditioned response as measured by self-reported fear, SCR, and activation of the right INS. Following the acquisition of differential fear, fear extinction learning is commonly used for inhibiting the fear conditioned association (Milad & Quirk, 2012). Fear extinction learning involves repeated presentations of the CS+ in the absence of the US such that a novel safety associated memory is formed, which leads to a reduction or suppression of the differential CR. Many imagery-based clinical interventions such as imaginal exposure (Abramowitz, 1996; Hecker, 1990; Powers et al., 2010; Turner et al., 1994) are premised on such extinction theorizing.

Though limited in number, existing research has provided supporting evidence for the efficacy of imagery exposure in the down-regulation of differential fear conditioning via fear extinction (e.g., Agren et al., 2017; Reddan et al., 2018). Agren et al. (2017) found that exposure to either an imagined or perceptual CS+ during extinction led to comparable degrees of reduction in differential SCR. Reddan et al. (2018) reported reductions in both threat-relevant neural predictive pattern expression and SCR upon re-exposure to the conditioned auditory CS+ following imagined extinction. During this imagined extinction participants were asked to play the conditioned CS+ tones in their head. Koizumi et al. (2016) also managed to reduce conditioned fear responses as measured by SCR and neural activities using counter conditioning by pairing implicit imagery of the CS+ with monetary reward using neurofeedback. Notably, the above cited studies all relied on physiological indices of differential conditioning without assessments of the subjective experiences of fear. Although there are important methodological reasons for avoiding the simultaneous acquisition of some measures of differential conditioning that may confound each other (e.g., SCR and fear potentiated startle; Sjouwerman, Niehaus, Kuhn, & Lonsdorf, 2016), from the perspective of the multi-system conceptualization of emotion it is important to collected multiple measures when possible. Specifically, emotion is expressed through both the conscious feeling of fear and also the corresponding behavioral and physiological expressions (LeDoux & Pine, 2016). Over-reliance on any single response system will limit our understanding of the emotion being studied (Frijda, 1986; Gross, 2013; Jiang, Burleigh, & Greening, 2021; Lang, 1993; Larsen & Prizmic-Larsen, 2006; LeDoux & Pine, 2016). For instance, Mauss and colleagues (2005) reported only moderate correlations between self-reported descriptions of the emotional experience and the corresponding physiological responses. In addition, neuroimaging evidence suggests that subjective and physiological expressions of fear may be reflective of different brain and cognitive mechanisms (Taschereau-Dumouchel, Kawato, & Lau, 2019).

The current study aimed to evaluate whether visual mental imagery could produce fear extinction following the acquisition of differential fear conditioning to visual objects, in terms of both self-reported fear and SCR. To do so, we used a within-subject design in which two CS+s and one CS- underwent acquisition of differential conditioning (Acquisition), followed by in vivo, perceptual extinction of one of the CS+s (CS+V) and imagery extinction of the other (CS+I) during Extinction Phase 1. Both of the CS+s were then viewed again in Extinction Phase 2. Considering the ample evidence supporting the effectiveness of imagery-based clinical interventions on the reduction of negative affect (Heyes, Lau, & Holmes, 2013; Holmes & Mathews, 2010; Weßlau & Steil, 2014) and the above research findings on imagery in fear down-regulation (Agren et al., 2017; Koizumi et al., 2016; Reddan et al., 2018), our overall prediction was that conditioned fear responses (as measured by both self-report and SCR) to perceived visual stimuli would be reduced after repeated imaginary exposure to the same stimuli without the accompanying US. The accuracy of this overall prediction was evaluated on the basis of two specific experimental predictions. First, we hypothesized that imagining the CS+I during Extinction Phase 1 would elicit a significant CR compared to the CS-, as the fear association would generalize from viewing the CS+I during Acquisition to imagining the CS+I during Extinction Phase 1 (Greening et al., 2021). Second, if fear extinction learning via imagery exposure generalized back to viewing the CS+I, then we would observe significant and similar reductions in CRs associated with the CS+V and CS+I during early Extinction Phase 2 consistent with previous research (Agren et al., 2017; Reddan et al., 2018). Conversely, if imagery exposure did not generalize to the visual CS+I, we would predict a persistent differential CR for the CS+I in early Extinction Phase 2.

## 2. Method

### 2.1 Participants

A total of 61 Louisiana State University students taking undergraduate-level psychology courses were recruited and completed this study. Five participants were removed from the final analyses due to excessive noise in SCR throughout or a lack of SCR response to the US, i.e., the shock, leaving a final sample of 56 individuals (32 women, mean age of 19.28, *SD* = 1.18; demographic information is missing from 3 participants). Our sample size was based on the research findings of Reddan et al. (2018) and Agren et al. (2017). However, we also conducted separate Sensitivity Analyses from a Bayesian framework (Krypotos, Klugkist, & Engelhard, 2017) on the SCR data for each study phase to test the robustness of CS+I versus CS- difference (two-sided). This used the Cauchy distribution with scaling factors of 0.1, 0.3, 0.5, 0.7, 1, 1.5 to interrogated how the choice of prior affected the resulting Bayes Factor. This study was approved by the Institutional Review Board of the Louisiana State University. Written informed consent was given to each participant prior to the start of experiment sessions. All participants were reimbursed by way of research credits.

### 2.2 Stimuli

All stimuli were created and presented in MATLAB R2018a with the Psychophysics Toolbox extensions (Brainard, 1997; Kleiner et al., 2007; Pelli, 1997). Three capital letters were selected as the CSs: letters “J” and “H” were paired with the US functioning as two distinct CS+s and letter “F” was used as the CS- and it was never paired with the US. All stimuli were presented at the center of the screen in 4-by-5 grids drawn on a grey background (figure 1A). A single mild electric shock of 5-ms duration as the US was delivered to the distal phalanx of each participant’s ring finger and pinky on his/her non-dominant hand through attached electrodes using the Biopac MP-150 system and AcqKnowledge software (for 5 participants, the shock electrodes were placed on their index and middle fingers). Whenever the US was delivered it coterminated with the CS+. The intensity of the US was customized to each participant such that it was rated as “uncomfortable but not painful.” Shock intensity was checked and adjusted after the first half of Acquisition for each participant to avoid desensitization or sensitization to the US.

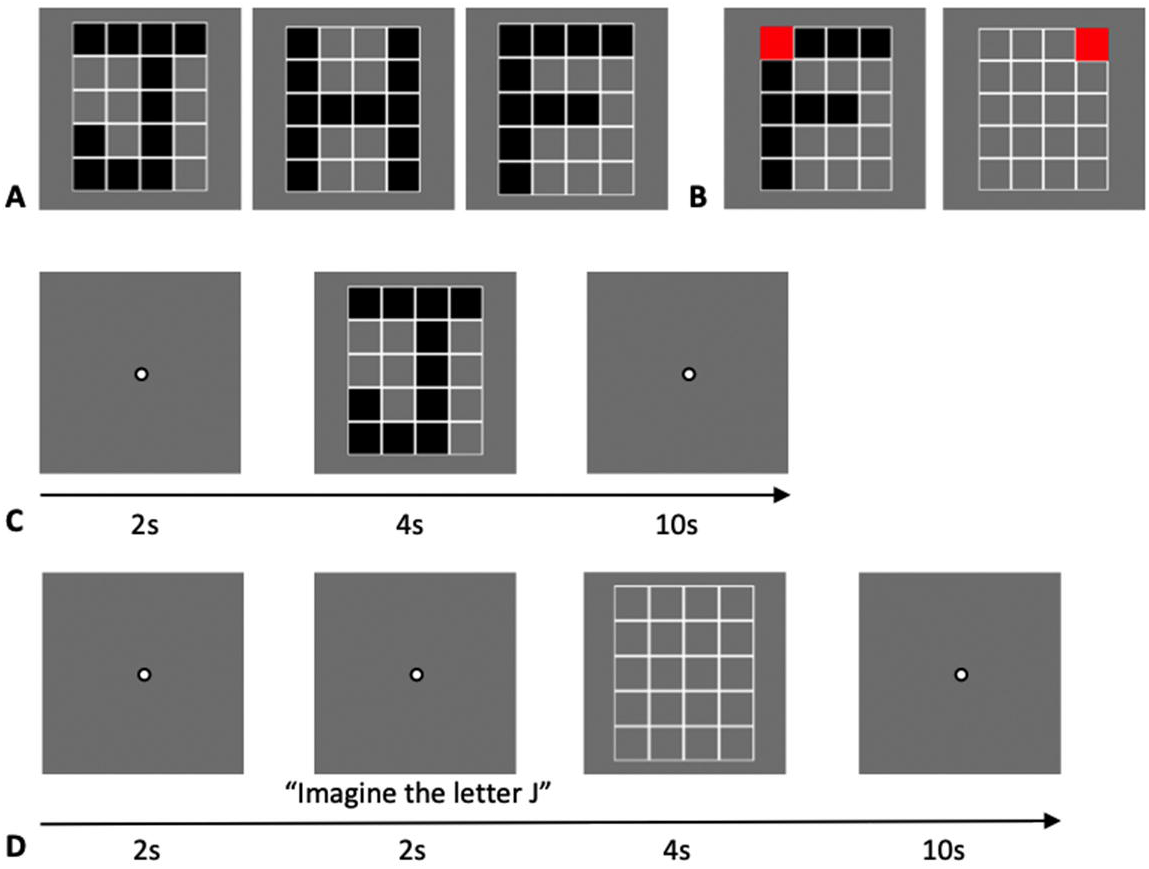

### 2.3 Procedures

Each participant first completed two training runs (12 trials/run) in which participants both viewed (figure 1C) or were instructed to imagine (figure 1D) the selected three letters. During each trial a probe appeared, which was a single square in the grid filled-in in red (figure 1B) and participants were instructed identify via button press whether the probe was on or off the letter viewed/imagined. Specifically, they were instructed to press 1 if they thought the probe was on the letter and press 2 if they thought the probe was off the letter. This probe task was added to the training phrase to initially facilitate participants’ attention to the task and ensure the instructions were understood regarding imagery. No shock was involved during this training phase of the experiment, which allows the trials in this phase to function as habituation trials for the CSs. However, no probes were present during the remaining phases of the experiment out of a concern that they could interfere with the differential conditioning (Carter et al., 2003).

During fear conditioning/Acquisition (12 trials/run x 4 runs; each CS letter, i.e., “J”, “H”, and “F”, was presented 16 times total), all trials were view trials (i.e., there was no mental imagery). The two CS+ letters were paired with shock on 50% of the trials (i.e., there were 8 trials per CS+ that were reinforced with the US) and the CS- was never paired with shock. Participants were told that “for 2 out of the 3 letters, there is a chance you may get shocked” before the beginning of conditioning. In a given run, the order of stimulus presentation was pseudorandomized in a way that the first and last trials of each run were always CS- trials (i.e., letter “F”). Additionally, each run contained two reinforced trials per CS+ (i.e., CS+V and CS+I, or the letters “J” and “H”). The second and third trials of each run were always reinforced shock trials for each of the CS+V and the CS+I (the order is randomized). The second reinforced CS+ trial for both the CS+V and the CS+I always occurred in the second half of a given acquisition run. In addition, no more than two consecutive trials were the same. Upon completion of Acquisition, participants proceeded to the Extinction Phase 1 (12 trials/run x 2 runs; each CS letter, i.e., “J”, “H”, and “F”, was presented 8 times total) in which shock was removed from CS+ presentations. One CS+ continued to be viewed (CS+V) and the other was imagined (CS+I) for the entire phase. The assignment of which CS+ letter (i.e., letters “J” and “H”) was the CS+V versus the CS+I was counterbalanced across participants. The CS- (i.e., letter “F”) was always viewed during Extinction Phase 1. The last phase of the experiment was Extinction Phase 2 (12 trials/run x 2 runs; each CS letter, i.e., “J”, “H”, and “F”, was presented 8 times total), in which all stimuli were viewed without the presence of electric shocks. The order of stimulus presentation for both Extinction Phase 1 and Extinction Phase 2 was pseudorandomized with no more than two of the same CS trials in a row. Details of the trial schedule for each phase are listed in table 1.

### 2.4 Self-Report Measures

After Acquisition, Extinction Phase 1 and 2, participants were asked to complete a series of self-report questionnaires on their level of fear towards receiving a shock, and on their perceived likelihood of being shocked for each CS. They were instructed to report how much they feared receiving a shock using a 7-point Likert scale (between 1 = “Not at All” and 7 = “Very Much So”). A 10-point Likert scale (between 0% and 100% with intervals of 10%) was provided for reporting estimations of shock likelihood.

### 2.5 Skin Conductance Response

During each trial, participants’ SCRs were collected and sampled at 2000 Hz. SCR analysis was carried out in MATLAB R2018a (Version 9.4). A first-order Butterworth bandpass filter was applied during preprocessing with cut-off frequencies of .01 and 5 Hz (Bach, Flandin, Friston, & Dolan, 2010). The time series were then down-sampled to 100 Hz. Based on previous research (Grégoire & Greening, 2019; Schiller, Kanen, LeDoux, Monfils, & Phelps, 2013; Vervliet, Vansteenwegen, Baeyens, Hermans, & Eelen, 2005) SCRs to the CSs were calculated by subtracting baseline (1 second before stimulus onset) from peak amplitude (during 1-3.995 seconds after stimulus onset). Peaks were zeroed if their distance from the baseline was smaller than .02 μS. The difference scores were then square root transformed. Consistent with previous research (Grégoire & Greening, 2019; Reddan et al., 2018; Schiller et al., 2010), we only included unreinforced CS+ trials (i.e., trials without US) in subsequent analyses. Lastly, the first and last trials of the Acquisition run were always CS- trials, which were excluded from the analyses to avoid the potential confounding orienting (Schiller et al., 2010) or “novelty” effect (Lim & Pessoa, 2008) that can occur on the first trial of a run, and to ensure that there was an equal number of CS+ and CS- trials included in the analyses (Grégoire & Greening, 2019, 2020; Lonsdorf et al., 2017).

### 2.6 Data Analysis

Based on recommendations by Lonsdorf et al. (2017), in order to maintain the most generalizable sample possible we only excluded participants from the analysis who displayed no clear SCR response to the US or whose overall SCR data was of poor quality even after filtering and down-sampling. No outlier removal was undertaken. Mean values of square-rooted SCR were calculated for each CS types (i.e., CS+I, CS+V, and CS−) separated into Acquisition, early Extinction Phase 1 (first half of trials), late Extinction Phase 1 (second half of trials), early Extinction Phase 2 (first half of trials), and late Extinction Phase 2 (late half of trials). Selfreported fear and shock estimations were collected at the end of each study phase and thus were not broken down into early and late responses for each of the three CSs. Repeated measures analyses of variance (ANOVAs) were conducted with Greenhouse–Geisser correction for non-sphericity when needed. Post hoc paired *t*-tests without corrections were conducted when significant interactions or main effects were found (Saville, 1990). Specific analyses are described in the results section.

## 3. Results

### 3.1 Acquisition

#### SCR

Acquisition was analyzed using a repeated-measures ANOVA with CS type (i.e., CS+I, CS+V, and CS-) as the within-subject factor on mean SCR during Acquisition (figure 2). There was a main effect of CS type, *F*(2, 110) = 3.80, *p* = .025, *η*^2^ = .06. Post hoc *t*-tests revealed that while there was a significant difference between the CS+I (*M* = .23, *SD* = .21) and the CS- (*M* = .17, *SD* = .14), *t*(55) = 2.59, *p* = .012, *d* = .35; the CS+V (*M* = .20, *SD* = .19) did not differ from the CS- (*M* = .17, *SD* = .14), *t*(55) = 1.40, *p* = .167. Moreover, the two CS+s (i.e., CS+I and CS+V) did not differ from each other, *t*(55) = 1.46, *p* = .151. To test the robustness of CS+I and CS- difference, a Sensitivity Analysis from a Bayesian framework (Krypotos et al., 2017) was conducted contrasting the CS+I and CS- (two-sided). The resulting Bayes Factors (BF_10_) were: BF_10_(0.1) = 3.357, BF_10_(0.3) = 4.178, BF_10_(0.5) = 3.658, BF_10_(0.7) = 3.062, BF_10_(1) = 2.381, BF_10_(1.5) = 1.694, respectively, indicating moderate to anecdotal evidence for accepting the alternative hypothesis. Together, based on the mean SCR, CS+I but not CS+V acquired differential fear at the group-level. Thus, we did not include the CS+V in the subsequent analyses in the extinction phases with respect to the SCR data.

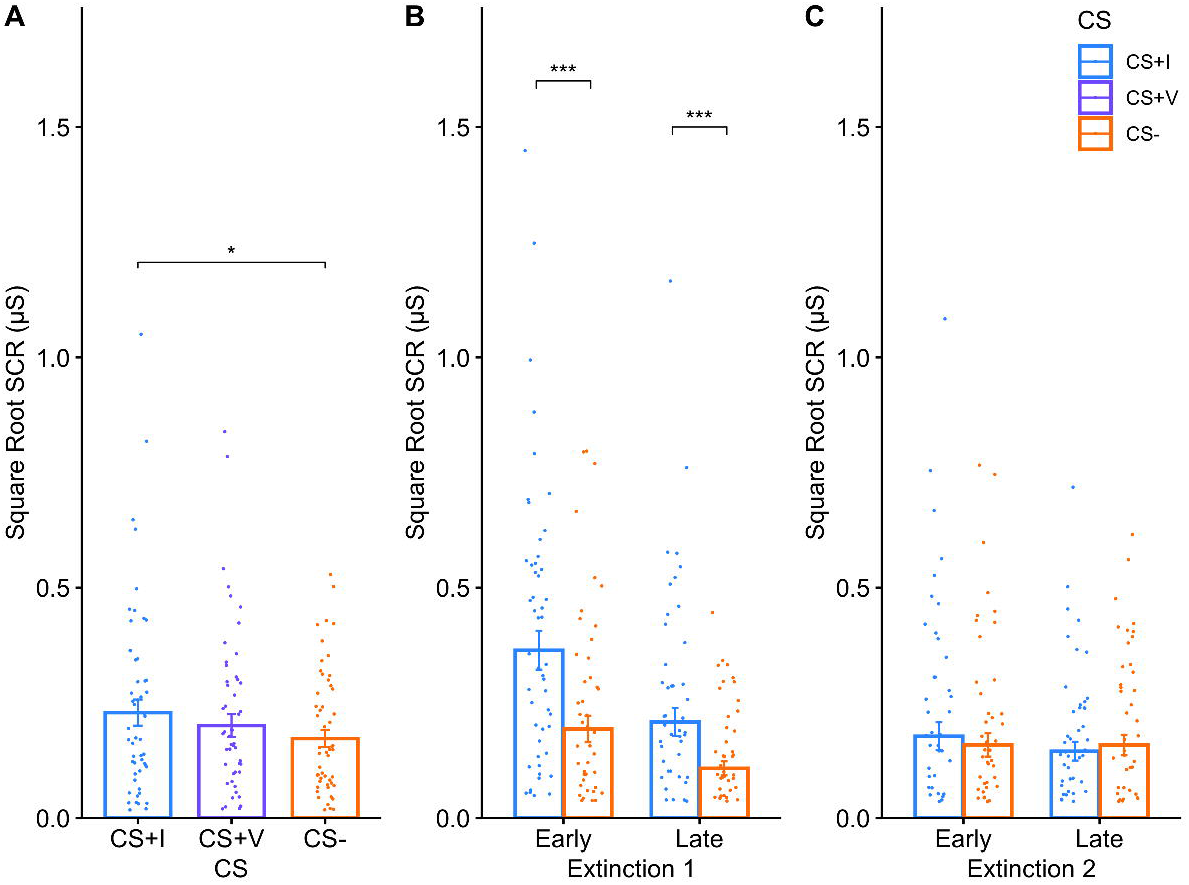

#### Self-Report

The same repeated ANOVA was applied to self-reported fear (figure 3A) and shock estimations (figure 3B). In contrast to SCR, these two self-report measures revealed differential conditioning of both CS+s. Specifically, for self-reported fear there was a main effect of CS-type, *F*(1.38, 76) = 32.59, *p* < .0001, *η*^2^ = .37. Post hoc *t*-tests indicated that participants reported higher fear ratings for both the CS+I (*M* = 4.20, *SD* = 1.63) and the CS+V (*M* = 4.23, *SD* = 1.67) compared to the CS- (*M* = 2.14, *SD* = 1.43), with respective statistics of *t*(55) = 6.61, *p* < .0001, *d* = .88, and *t*(55) = 7.13, *p* < .0001, *d* = .95. The two CS+s did not differ, *t*(55) = −.12, *p* = .901. As for shock estimation, a main effect of CS type was also found, *F*(1.71, 94.22) = 59.95, *p* < .0001, *η*^2^ = .52. Post hoc *t*-tests showed that participants reported higher likelihood of shock for both the CS+I (*M* = 49.82, *SD* = 17.11) and the CS+V (*M* = 49.64, *SD* = 18.39) compared to the CS- (*M* = 17.86, *SD* = 20.95): CS+I vs CS-, *t*(55) = 8.45, *p* < .0001, *d* = 1.13; CS+V vs CS-, *t*(55) = 8.07, *p* < .0001, *d* = 1.08). The two CS+s, again, did not differ, *t*(55) = .06, *p* = .951.

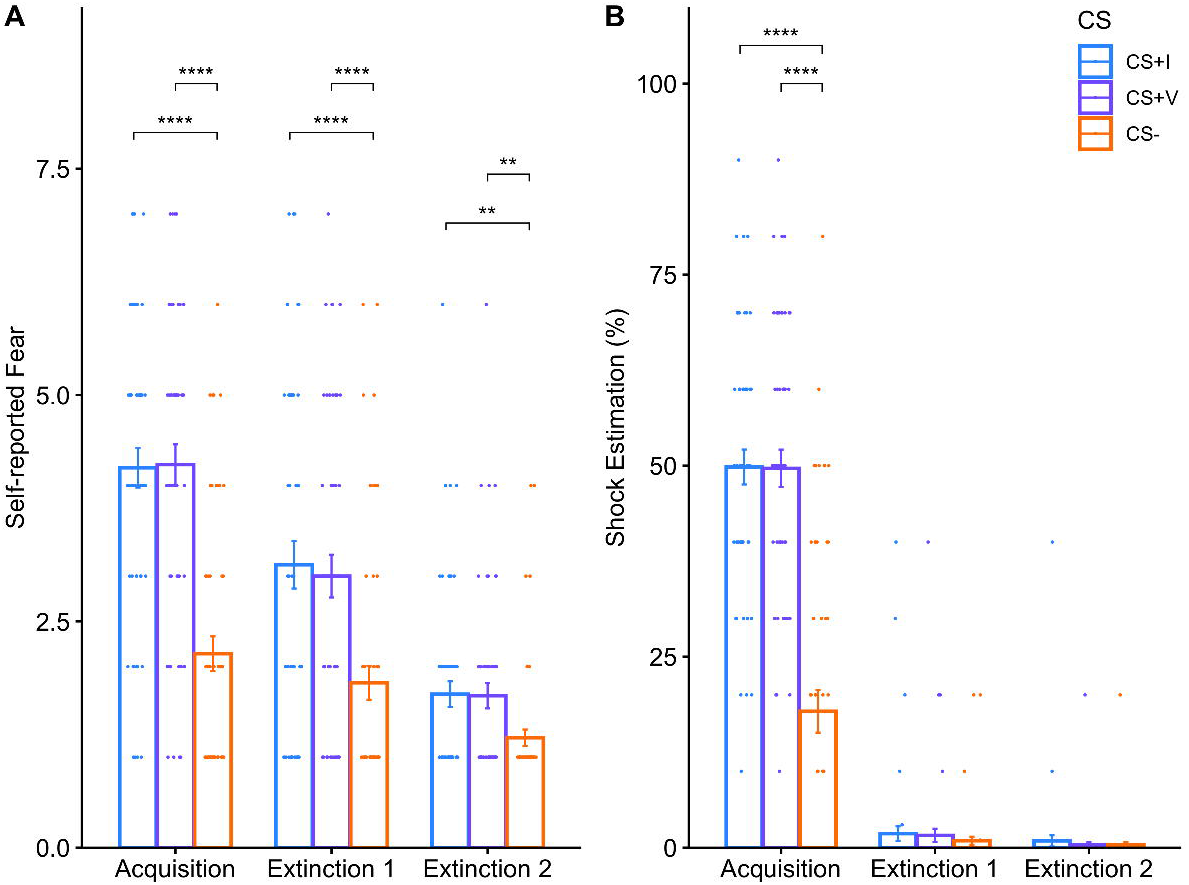

### 3.2 Extinction Phase 1: Differential Fear Generalization to CS+ Imagery

#### SCR

During Extinction Phase 1, participants imagined the CS+I and continued to view the CS-, and the US was never delivered. A 2×2 ANOVA with CS-type (i.e., CS+I, CS-) and timing (i.e., early trials, late trials) as the within subject factors were conducted on SCR data during Extinction Phase 1 (figure 2B). This analysis revealed significant main effects for both CS type, *F*(1, 55) = 21.12, *p* < .0001, *η_p_*^2^ = .28, and timing, *F*(1, 55) = 41.45, *p* < .0001, *η_p_*^2^ = .43. There was no CS type x timing interaction, *F*(1, 55) = 2.91, *p* = .094. Post hoc *t*-tests indicated that the CS+I was significantly different from the CS- during both early (CS+I: *M* = .38, *SD* = .32; CS-: *M* = .19, *SD* = .20), *t*(55) = 4.03, *p* < .001, *d* = .54; and late (CS+I: *M* = .22, *SD* = .24; CS-: *M* = .11, *SD* = .12), *t*(55) = 3.55, *p* < .001, *d* = .47 trials of Extinction Phase 1. Regarding the robustness of the CS+I versus CS- difference, the resulting Bayes Factors (BF_10_) for early Extinction Phase 1 were: BF_10_(0.1) = 64.462, BF_10_(0.3) = 132.152, BF_10_(0.5) = 144.646, BF_10_(0.7) = 136.552, BF_10_(1) = 116.669, BF_10_(1.5) = 88.786, respectively, indicating very strong to extreme evidence for accepting the alternative hypothesis. For late Extinction Phase 1, the calculated Bayes Factors (BF_10_) were: BF_10_(0.1) = 20.209, BF_10_(0.3) = 36.588, BF_10_(0.5) = 37.432, BF_10_(0.7) = 33.971, BF_10_(1) = 28.095, BF_10_(1.5) = 20.871, indicating strong to very strong evidence for accepting the alternative hypothesis.

Both the CS+I and the CS- had greater SCR during early compared to their respective late trials of Extinction Phase 1 (CS+I: *t*(55) = 5.53, *p* < .0001, *d* = .74; CS-: *t*(55) = 3.10, *p* = .003, *d* = .41). The robustness of the difference between early and late trials for the CS+I was indicative of extreme evidence for the alternative hypothesis, the Bayes Factors (BF_10_) were all greater than 1000. As for the CS-, the reported Bayes Factors (BF_10_) were: BF_10_(0.1) = 7.857, BF_10_(0.3) = 12.149, BF_10_(0.5) = 11.572, BF_10_(0.7) = 10.105, BF_10_(1) = 8.109, BF_10_(1.5) = 5.896, indicating moderate to strong evidence supporting the alternative hypothesis. As noted previously, the CS+V was not included in SCR analyses for Extinction Phase 1 and 2. However, a graph containing all three CSs separated into early and late Extinction Phase 1 and 2 is provided in supplementary figure S1.

#### Self-Report

Results of a one-way repeated measures ANOVA with CS-type (i.e., CS+I, CS+V, and CS-) as the within-subject factor on self-reported fear (figure 3A) also supported generalization of fear to imagery with a main effect of CS-type, *F*(1.75, 96.49) = 15.88, *p* < .0001, *η_p_*^2^ = .22. Based on post hoc *t*-tests, CS+I (*M* = 3.28, *SD* = 1.96) and CS+V (*M* = 3.16, *SD* = 1.77) were both rated significantly higher than CS- (*M* = 1.92, *SD* = 1.45), with *t*(55) = 4.36, *p* < .0001, *d* = .58, and *t*(55) = 5.10, *p* < .0001, *d* = .68, respectively. The two CS+s did not differ from each other, *t*(55) = .55, *p* = .588. The same one-way repeated measures ANOVA applied to shock estimations indicated that all CS’s (i.e., CS+I, CS+V, and CS-) were reported to be paired with shock close to 0% of the time during Extinction Phase 1 (CS+I: *M* = 1.84, *SD* = 7.16; CS+V: *M* = 1.61, *SD* = 6.54; CS-: *M* = .91, *SD* = 3.94) with no effect of CS type, *F*(1.47, 81.02) = 1.15, *p* = .308.

### 3.3 Extinction Phase 2: Imagery Exposure Led to Differential Fear Extinction Learning

#### SCR

During Extinction Phase 2, participants now viewed both the CS+I and the CS-, and as with Extinction Phase 1 the US was never delivered. We predicted that if mental imagery of CS+I was effective then we would observe no main effect of CS type. As predicted, a 2×2 ANOVA with CS-type (i.e., CS+I, CS-) and timing (i.e., early and late extinction) as the within subject factors (figure 2C) resulted in no interaction (*F*(1, 55) = .52, *p* = .476) nor main effects (CS type: *F*(1, 55) = .02, *p* = .882; timing: *F*(1, 55) = .49, *p* = .485). This suggests that CS+I and CS- did not differ during either early (CS+I: *M* = .18, *SD* = .23; CS-: *M* = .16, *SD* = .19) or late (CS+I: *M* = .15, *SD* = .15; CS-: *M* = .16, *SD* = .17) Extinction Phase 2. Regarding the robustness of this lack of difference between the CS+I and the CS- in the early Extinction Phase 2, the resulting Bayes Factors (BF_01_) were: BF_01_(0.1) = 1.534, BF_01_(0.3) = 2.813, BF_01_(0.5) = 4.231, BF_01_(0.7) = 5.711, BF_01_(1) = 7.982, BF_01_(1.5) = 11.823, indicating anecdotal to strong evidence in support of the null hypothesis. For late Extinction Phase 2, the Bayes Factors (BF_01_) were: BF_01_(0.1) = 1.575, BF_01_(0.3) = 2.945, BF_01_(0.5) = 4.459, BF_01_(0.7) = 6.035, BF_01_(1) = 8.45, BF_01_(1.5) = 12.529, again, providing anecdotal to strong evidence supporting the null hypothesis.

#### Self-report

Regarding self-reported fear, a one-way repeated measures ANOVA with CS-type (i.e., CS+I, CS+V, and CS-) as the within-subject factor (figure 3A) revealed a main effect of CS type, *F*(1.75, 96.24) = 14.25, *p* < .0001, *η_p_*^2^ = .21. Importantly, a post hoc *t*-test revealed no significant difference in self-reported fear for the CS+I (*M* = 1.70, *SD* = 1.06) versus the CS+V (*M* = 1.68, *SD* = 1.05), *t*(55) = .10, *p* = .924. Additional post hoc *t*-tests revealed higher fear ratings for both the CS+I and the CS+V than the CS- (*M* = 1.21, *SD* = .68), with *t*(55) = 2.89, *p* = .006, *d* = .39, and *t*(55) = 3.00, *p* = .004, *d* = .40, respectively. Although this result may appear to suggest that self-reported fear was not extinguished, in the following section we carried out an additional analysis across the 3 main experimental phases to demonstrate that the magnitude of self-reported differential fear was reduced via both visual and imagery extinction. Regarding self-reported shock estimation, accurate estimations were reported (figure 3B) with respect to Extinction Phase 2 with close to 0% estimations for the CS+I (*M* = .89, *SD* = 5.49), the CS+V (*M* = .38, *SD* = 2.67), and the CS- (*M* = .36, *SD* = 2.67) with no main effect of CS type, *F*(1, 55) = 1.83, *p* = .182.

### 3.4 Self-Reported Fear Across Acquisition, Extinction Phase 1 and Extinction Phase 2

In order to evaluate the magnitude of differential fear conditioning as measured by selfreport across the three main experimental phases we conducted a 3×2 ANOVA on the difference scores of self-reported fear with study phase (i.e., Acquisition, Extinction Phase 1, Extinction Phase 2) and CS contrast (i.e., CS+I minus CS-, CS+V minus CS-) as the within-subject factors. This revealed only a main effect of study phase, *F*(1.71, 94.27) = 17.84, *p* < .0001, *η_p_*^2^ = .24. Notably, there was neither a main effect of CS contrast, *F*(1, 55) = .13, *p* = 721, nor interaction, *F*(1.45, 79.87) = .23, *p* = .724. As there was no main effect of CS contrast and no interaction, the contrast scores were averaged together (i.e., producing one aggregate measure of the CS+ vs CS- difference) and compared between study phases. Post-hoc *t*-tests revealed significant reductions from Acquisition (*M* = 2.07, *SD* = 2.48) to Extinction Phase 1 (*M* = 1.24, *SD* = 1.81), *t*(55) = 2.83, *p* = .007, *d* = .38; and from Extinction Phase 1 to Extinction Phase 2 (*M* = .47, *SD* = .77), *t*(55) = 3.73, *p* = .000, *d* = .50. These results suggest that the magnitude of differential fear conditioning (CS+ minus CS-) as measured with self-reported fear significantly decreased throughout extinction (i.e., Extinction Phase 1 and 2). Moreover, the degree of reduction in differential self-reported fear was similar regardless of whether in vivo or imagery exposure was employed during Extinction Phase 1.

## 4. Discussion

Mental imagery has long played an important part in our understanding and treatments of various affective conditions. The assumption is that imagery performs like its perceptual counterpart and can exert regulatory effects on emotion in its place. While efforts have been made in the theorizing and assessment of imagery-based emotion regulatory strategies, experimental examination is lacking in comparison. The current investigation sought to determine if visual mental imagery could contribute to fear extinction as measured by both physiology (i.e., SCR) and self-reported fear, as this has not been done in the current literature (Agren et al., 2017; Reddan et al., 2018). First, we found that conditioned fear acquired to visual stimuli generalized to visual imagery as indicated by elevated SCR and self-reported fear. Second, following repeated visual imagery of the CS+I, there was no significant differential SCR upon re-exposure to the CS+I compared to the CS-. In addition, based on self-reported fear, imagery-based and perceptual extinction led to similar degrees of reductions in the differential fear conditioned response, consistent with the SCR-based findings of Agren et al. (2017).

The main research question of the current investigation was whether imagery exposure to the CS+ leads to a generalized fear reduction to the perceptual (i.e., visual) CS+. Therefore, our observation of significantly greater SCR for the CS+I versus the CS- during acquisition allows us to make valid inferences regarding mental imagery’s impact on extinction despite the lack of clear differential conditioning between the CS+V and the CS-. Our first observation supporting the utility of imagery exposure during extinction learning comes from the observation of the generalization of conditioned fear from perception to imagery during Extinction Phase 1 in terms of both SCR and self-reported fear. An alternative possibility is that the elevated SCR when imagining the CS+I could be the result of the act of imagery, which may require more effort than simply viewing in the case of the CS-. The initial purpose of including the CS+V during Extinction Phase 1 was to help evaluate this possibility. On the other hand, previous research on the generalization of differential fear conditioning from visual to imagined objects did not observe evidence of an elevated SCR response to mental imagery compared to viewing, though they did observe a significant SCR difference between the CS+ versus the CS- (Greening et al., 2021). Another study reported that while the conditioned response, as measured by the SCR difference between CS+ and CS-, was initially larger in the imaginal extinction group, the imagery and in vivo/perceptual extinction groups were comparable by late extinction (Agren et al., 2017). Based on these findings, it is unlikely that the SCR difference can be completely accounted for by the difference between viewing versus visual imagery per se, rather than being an expression of the generalized conditioned fear response.

The second piece of evidence supporting the use of mental imagery for fear extinction learning comes from Extinction Phase 2. Specifically, even during the early trials (i.e., the first four trials per condition) the CS+I no longer exhibited a greater SCR compared to the CS-. Had imagery fear extinction not affected the fear conditioned association between the perceptual in vivo CS+I and the CS- one would have predicted a persistent differential conditioning upon reexposure to the in vivo CS+I during Extinction Phase 2, which was not observed. Unlike SCR, differential fear measured as self-reported fear did persist to some degree throughout extinction. However, this differential was significantly reduced from Acquisition to Extinction Phase 1 and from Extinction Phase 1 to Extinction Phase 2 irrespective of the type of CS+ (i.e., for both CS+I – CS- and CS+V – CS-). This is in line with previous research on self-reported fear (Lau et al., 2008; Shechner et al., 2015), and may reflect that individuals maintain stable declarative knowledge regarding the difference between the CS+s and the CS- even after extinction.

While the present findings provide evidence that mental imagery can facilitate extinction learning, there are at least two potential mechanistic explanations underpinning the observed effects. The first possibility is that mental imagery operates via depictive mechanisms (Lewis et al., 2013) such that during extinction, the imagery of the CS+ acts as a substitute for the in vivo (i.e., visual) stimulus (e.g., Greening et al., 2021). Alternatively, our results could be the result of propositional mechanisms, e.g., the controlled reasoning that viewing the CS+ is no longer accompanied by the US since imagining the CS+ was not (Mitchell, De Houwer, & Lovibond, 2009). Future investigations are needed to further evaluate the underlying cognitive mechanisms of fear extinction during affective mental imagery, for example, by controlling for the potential confound of verbalization as has been done in previous research on mental imagery and emotion (Holmes & Mathews, 2005). Also see Mertens et al. (2020) for additional considerations when designing control conditions. Not only will such future investigations better our understanding of the operations of affective mental imagery, they would also be of great clinical significance (Andrade, May, & Kavanagh, 2012; Hirsch & Holmes, 2007).

Unexpectedly, only one of the two CS+s (i.e., the CS+I) was found to exhibit threat related differential physiological reactivity (i.e., greater SCR for the CS+I vs. the CS-) following the Acquisition phase in the current study. Considering that the letter selections of the CS+I and the CS+V were counterbalanced across participants, this could not be attributed to differences between the letter stimuli of “J” and “H”. One possibility is that the nature of the stimuli and the use of two CS+s in a within-subject designed made the stimulus discriminability for the purposes of differential conditioning difficult. Another potential contributor could be the length of the Acquisition phase. While a shock check was conducted during Acquisition, participants still exhibited habituation to the US towards the second half of Acquisition based on visual examination of the trial-by-trial SCR (see supplementary figure S2). Additionally, compared to evolutionarily fear-relevant objects (e.g., images of snakes, human faces), stimuli lacking in evolutionary fear-relevance, such as the letter stimuli used in the current study, undergo fear conditioning less readily (Öhman & Mineka, 2001). Future research could consider using more fear-relevant objects (e.g., snakes or spiders), shorter acquisition procedures, or a between-group design to avoid requiring two CS+s (e.g., Agren et al., 2017; Reddan et al., 2018). The inconsistency observed in the current study could also be a manifestation of the noise and measurement error in SCR.

Altogether, the present results from both SCR and self-reported fear suggest that repeated exposure to the mental image of an in vivo CS+ can facilitate extinction learning. Both the complementary findings and inferential similarities revealed between the subjective and physiological measures also demonstrate the importance of collecting multiple outcome measures based on a multi-component view of emotion (Frijda, 1986; Gross, 2013; Lang, 1993; Larsen & Prizmic-Larsen, 2006; LeDoux & Pine, 2016). The current study along with the existing experimental literature (Agren et al., 2017; Reddan et al., 2018) provide preliminary support for the potential clinical application of extinction-based imagery treatments in fear-related affective conditions.

