## Supplemental for "Psychophysiological Evidence for Fear Extinction Learning via Mental Imagery"

**Supplemental File**

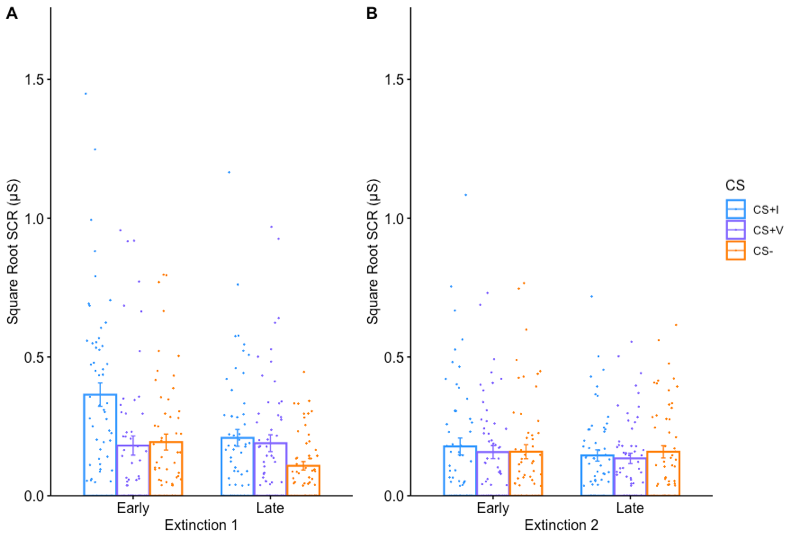

Figure S1. Mean square-rooted SCR for each CS type during early and late Extinction Phase 1 (A) and Extinction Phase 2 (B). Error bars show ± 1 standard error. Each dot represents one subject.

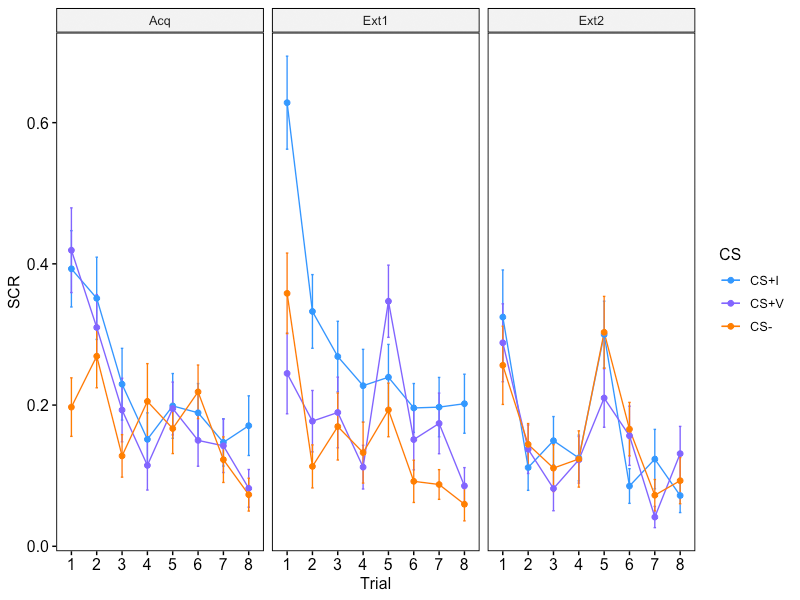

Figure S2. Trial-by-trial square-rooted SCR for each CS type across all three phases. Error bars show ± 1 standard error.
